# Distinct tasks engage a shared neural subspace in human hippocampus and anterior cingulate cortex

**DOI:** 10.64898/2026.04.24.720703

**Authors:** Hung-Yun Lu, Assia Chericoni, Melissa Franch, Kalman A. Katlowitz, Elizabeth Mickiewicz, Danika L. Paulo, Andrew J. Watrous, Eleonora Bartoli, Nicole R. Provenza, Sameer A. Sheth, Benjamin Y. Hayden, Jay A. Hennig

## Abstract

A hallmark of human cognitive flexibility is the ability to perform a wide range of unrelated tasks. While much is known about how neural circuits support individual tasks, less is understood about how these circuits support multiple tasks. Are neural representations largely task-specific, or do they retain a shared structure across distinct behaviors? To investigate this question, we recorded neural population activity from the hippocampus and anterior cingulate cortex, two regions linked to generalization and cognitive control, in eleven human patients performing three distinct tasks. Using dimensionality reduction, we estimated the neural subspace associated with each task and compared subspace geometry across tasks. We found that task-related subspaces were not independent: across tasks, approximately half of the subspace dimensions were shared, and these were primarily the dimensions containing most of the shared neural covariance. These findings indicate that neural population activity in these regions is not purely task-specific. This suggests a stable, low-dimensional circuit structure may persist even across unrelated behaviors, potentially providing a common substrate for flexible cognition.

## Introduction

The human brain has the remarkable ability to nimbly shift between tasks. This skill is important for daily life and its impairment is diagnostic of many psychiatric diseases, including depression and schizophrenia [1–3]. While most neuroscience research has focused on understanding how neural circuits support single tasks, much less attention has been paid to understanding how the same neural circuits reconfigure across different tasks. In particular, it is unclear to what extent neural circuits are specialized for individual tasks versus generally useful for multiple tasks. Answering this question is critical for understanding the broader human capacity to respond adaptively in a variety of contexts, and may be key for developing brain-inspired models of artificial general intelligence.

A promising approach to understanding how the brain supports multiple tasks is to examine neural activity at the population level [4–6]. The collective activity of neural populations occupies a low-dimensional subspace [7,8], commonly referred to as a neural manifold [9–11]. The manifold perspective has led to novel insights about function throughout the brain, including sensory cortex [12–14], motor cortex [15–18], frontal cortex [19–22], and hippocampus [23–26]. From a mechanistic perspective, the manifold is thought to arise from underlying biological properties such as circuit connectivity [11, 15, 27, 28]. From a functional perspective, the manifold is thought to reflect task-relevant computations (e.g., [6, 9, 10, 14, 29–31]). For example, features such as the dimensionality of the manifold may reflect stimulus or task complexity [9, 10, 29, 32]. The neural manifold thus provides a window into how the brain performs task-relevant computations.

Studies of both non-human animals and artificial agents have begun to shed light on how neural manifolds relate to the performance of multiple tasks, following extensive training on a collection of tasks [16, 24, 30, 33–39]. However, in contrast to non-human animals and artificial agents, humans are able to rapidly learn and perform multiple tasks without extensive training. What role do neural manifolds have in shaping this rapid flexibility? This question, which to our knowledge has not been previously investigated, highlights an underlying tension between the idea that the neural manifold is somehow both constrained by underlying circuitry and yet also reflective of concurrent task demands.

Here we consider two hypotheses for how neural manifolds, or *subspaces*, might support multiple tasks. One possibility is that neural subspaces are inherently adaptable to the task at hand [28, 30, 37, 40]. Specifically, a neural circuit may be able to rapidly reconfigure its subspace to match the computations needed for each given task. Under this hypothesis, task-specific computations are enabled by task-specific subspaces (Fig 1A). These different subspaces could arise if different tasks recruit entirely separate brain regions, disjoint sets of neurons in the same region (Fig 1C), or even separate functional subspaces within the same set of neurons (Fig 1D). Alternatively, on short timescales, neural circuits may not be able to rapidly reconfigure their neural subspaces due to underlying biological constraints such as circuit connectivity [15, 41–43]. As a result, neural circuits may execute different tasks by reusing (or “recycling” [44]) a shared subspace (Fig 1B). Under this hypothesis, task-specific activity unfolds within a fixed subspace that is used for multiple tasks [16, 43, 45–48]. In other words, while the neural computations performed by an area may change across tasks, the subspace itself would remain largely unchanged. For example, as depicted in Fig 1B, the neural trajectories across tasks may be task-specific, while the subspaces within which each trajectory evolves are nearly identical. Testing these hypotheses, by examining how neural population activity changes across multiple tasks, could potentially offer key insights into the neural mechanisms underlying human flexibility.

**Fig 1.**
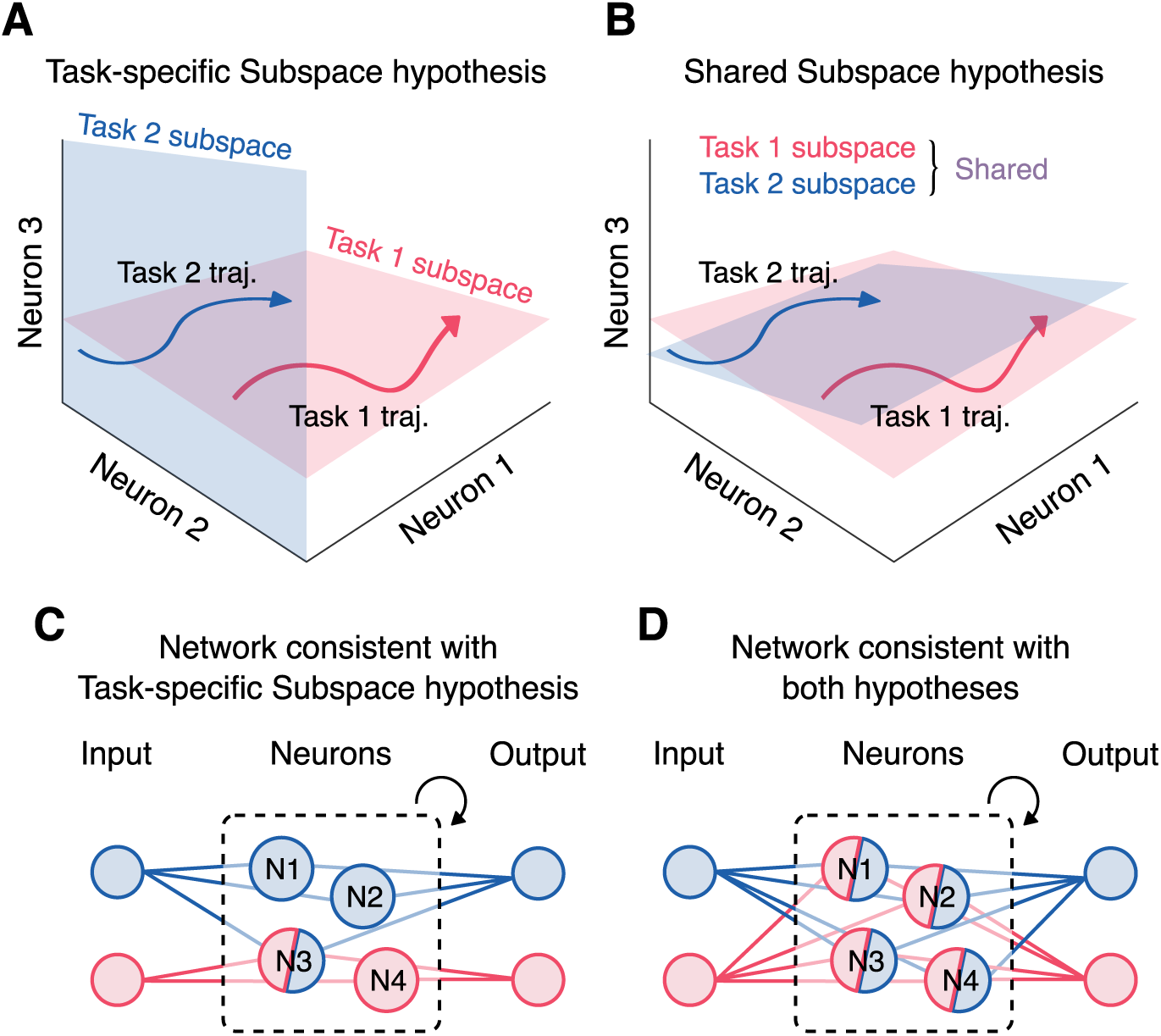
Two hypotheses for how neural subspaces may relate across tasks. **(A)**. In the Task-specific Subspaces hypothesis, the neural trajectories observed in different tasks (e.g., Task 1 and Task 2 trajectories shown in red and blue, respectively) occur in subspaces that are task-specific. **(B)**. In the Shared Subspace hypothesis, the neural trajectories observed in different tasks occur in subspaces that are highly overlapping, or “shared”. **(C)**. Example of a network that would give rise to task-specific subspaces, for two tasks with distinct inputs and outputs. Neurons active during Task 1 are colored red, neurons active during Task 2 are colored blue, and neurons colored red and blue are active during both tasks. In this network, most neurons are active during only a single task. **(D)**. Example of a network that could give rise to either task-specific or shared subspaces, with same conventions as panel C. Here, all neurons are active during both tasks.

Here we examined whether the brain uses task-specific or shared neural subspaces across multiple unrelated tasks. We were especially interested in the hippocampus (HPC), a region associated with generalization and cognitive mapping [49, 50], as well as in the anterior cingulate cortex (ACC), a brain region closely implicated in task-switching and executive control [51]. We recorded from neural population activity in both regions in eleven human patients as they performed three distinct tasks. For each patient, we used dimensionality reduction to characterize the neural subspace during each task. We then compared the resulting neural subspaces across tasks and found that they were much more similar than expected by chance, even though the tasks themselves involved distinct sensory, cognitive, and motor demands. We found that task-conditioned subspaces had substantial overlap across all pairs of tasks. While these subspaces were not perfectly aligned, the shared components of the subspaces contained the bulk of the neural covariability in each task. This indicates that, as humans perform multiple unrelated tasks, neural population activity in HPC and ACC largely occupies a shared neural subspace. These findings suggest that, across tasks, neural circuits maintain a stable, low-dimensional structure that may underlie flexible cognition.

## Results

We recruited eleven human patients undergoing neural monitoring for epilepsy. During the course of their stay, these patients performed at least two out of three distinct tasks (Fig 2A-C) while we continuously recorded spiking activity. In the Prey-pursuit task (*Pursuit*), patients used a joystick to pursue and capture continuously moving “prey” stimuli [26, 52] (Fig 2A). In the Podcast listening task (*Podcast*), native English-speaking patients listened to 47 minutes of English speech, taken from six monologues from The Moth podcast [53] (Fig 2B). In the Mental Rotations task (*Rotations*), patients were shown two block stimuli and responded with whether they were rotationally equivalent or mirror images (Fig 2C).

**Fig 2.**
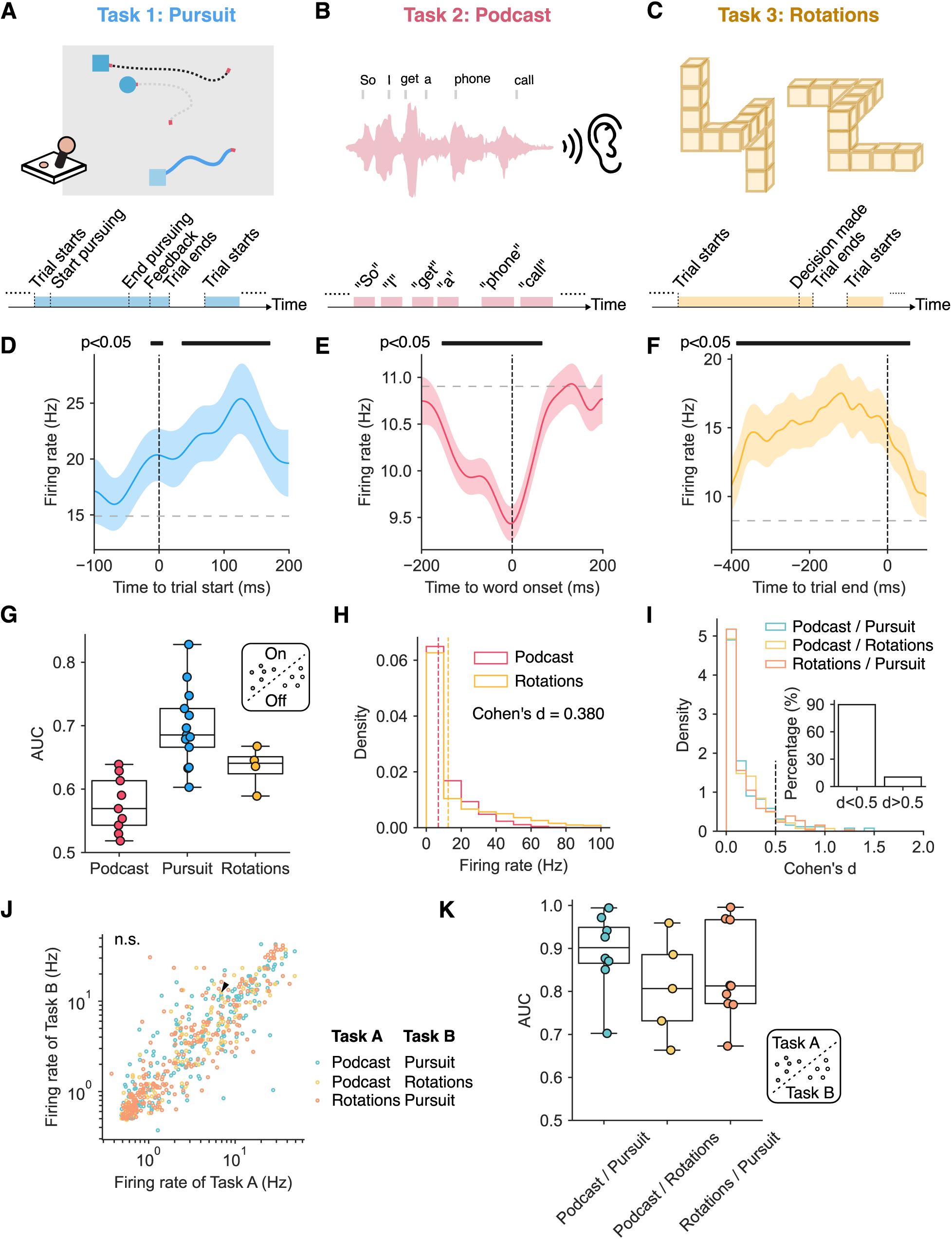
Neural units were tuned to three unrelated tasks. **(A)**. In the Prey-pursuit (“Pursuit”) task [26], patients captured prey (squares) by controlling the position of an avatar (circle) on a screen using a joystick. **(B)**. In the Podcast listening task, patients listened to six stories from The Moth podcast [53]. **(C)**. In the Mental rotations (“Rotations”) task, patients judged whether two block stimuli were rotationally equivalent or mirror images. **(D-F)**. Time-smoothed firing rates of representative units were averaged across trials from example Pursuit (panel D), Podcast (panel E), and Rotations (panel F) sessions. Gray horizontal lines indicate each unit’s mean firing rate across the full session. Colored lines indicate the trial-averaged event-aligned firing rates. Shaded regions indicate standard error of the means across trials. Two-tailed one-sample t-tests were used to assess whether firing rates at each time point differed significantly from the unit’s session mean, with false discovery rate (FDR) correction applied across time points. Time points where firing rates were significantly different from session mean (*p <* 0.05) were marked on top of each plot. **(G)**. The areas under the ROC curve of decoders trained to distinguish task status (“on” versus “off”) using normalized population firing rates. Each dot represents a single session. Whiskers show the minimum and maximum of the same group. **(H)**. The distributions of the firing rates from a representative unit recorded in Podcast and Rotations tasks. Dashed lines represent the means of the distributions. **(I)**. The distributions of Cohen’s *d* in different pairs of tasks. Inset: percentages of units that had small-to-moderate changes versus great changes in firing rates. **(J)**. The mean firing rates of units across different pairs of tasks. Each data point represents a unit. The black triangle indicates the example unit shown in panel H.**(K)**. The areas under the ROC curve of decoders trained to distinguish task identity (“task A” versus “task B”) using raw population firing rates. Each dot represents a pair of sessions. Whiskers show the minimum and maximum of all pairs of sessions.

Patients performed these tasks in various orders, and most tasks were performed within 30 hours of each other (Fig S1). In total, 11 patients completed 29 sessions across the three tasks (Fig S1; see Methods). Data from the Podcast and Pursuit tasks was previously analyzed in [26, 53]. Here, we asked how these same neurons participated across our three tasks.

### Neural activity was tuned to each task but differed across tasks

During these tasks we recorded spiking activity in HPC and ACC. While the activity on each channel may be composed of single- or multi-unit activity, we considered each channel as a single “neural unit,” allowing us to easily compare unit activity across tasks. We defined spike counts for each unit as the number of action potentials in 100 ms bins during each task (see Methods). We analyzed activity from units localized in HPC and ACC, resulting in around 37 ± 14 simultaneously recorded units per session (mean ± s.d., *N* = 29 sessions). We first asked whether neural activity was tuned to each task. We have shown in previous work on the same experiments that these units have task-specific encoding, including representations of spatial variables in the Pursuit task [26], and semantic content during the Podcast task [53]. These three tasks have rich structure, but they share no common task variables. To characterize task tuning in a uniform way across tasks, we defined “task-on” and “task-off” periods for each task (Fig 2A-C; see Methods). We found that firing rates were systematically modulated by these task events, including trial onset in the Pursuit task, word presentation in the Podcast task, and response periods in the Rotations task (representative units shown in Fig 2D–F). At the population level, we trained linear decoders to use neural population activity (defined as the instantaneous spike counts across all units) to predict task-on versus task-off periods. We quantified decoder performance using the area under the receiver operating characteristic curve (AUC). Across all sessions, decoder performance was significantly above chance levels (Fig 2G; permutation test, *N* = 1000, *p <* 0.001 for each session). Together, these results confirm that neural activity in HPC and ACC was strongly tuned to relevant task variables in each task.

We next asked how neural activity differed across tasks. At the single-unit level, in an example unit the firing rate changed significantly between the Podcast and Rotations tasks, suggesting that firing was task-specific. However, the effect size was relatively small (Fig 2H, Cohen’s *d* = 0.380). To assess whether this was typical across the population, we computed Cohen’s *d* for all units across task pairs and found that approximately 90% showed small-to-moderate changes in firing rates (Fig 2I, *d <* 0.5). Consistent with this, the mean firing rates across all units did not differ significantly between tasks (Fig 2J; two-tailed paired t-test, *T* (655) = 0.17, *p* = 0.866). At the population level, we trained linear decoders to use neural population activity to classify task identity between each pair of tasks. Decoding performance was well above chance levels across all task pairs (Fig 2K). Thus, while the majority of units were active across tasks, neural population activity also carried task-specific information sufficient to identify which task was being performed.

These results establish that neural activity in these experiments was modulated by each task while also differing systematically across tasks. Our results were more consistent with the network structure in Fig 1D because the majority of units were active across tasks. This sets the stage for our core question: whether population activity across tasks occupies task-specific or shared subspaces.

### Shared dimensionality and shared variance per unit were similar across tasks

To ask whether the neural population activity recorded across tasks was consistent with multiple task-specific subspaces or a single shared subspace, we took the following approach (Fig 3A). First, we identified a neural subspace during each task using the spike counts recorded from all simultaneously recorded units in a given patient. We will refer to each of these as a *task-conditioned subspace*. In the next section, we will ask whether the task-conditioned subspaces were the same or different across tasks.

**Fig 3.**
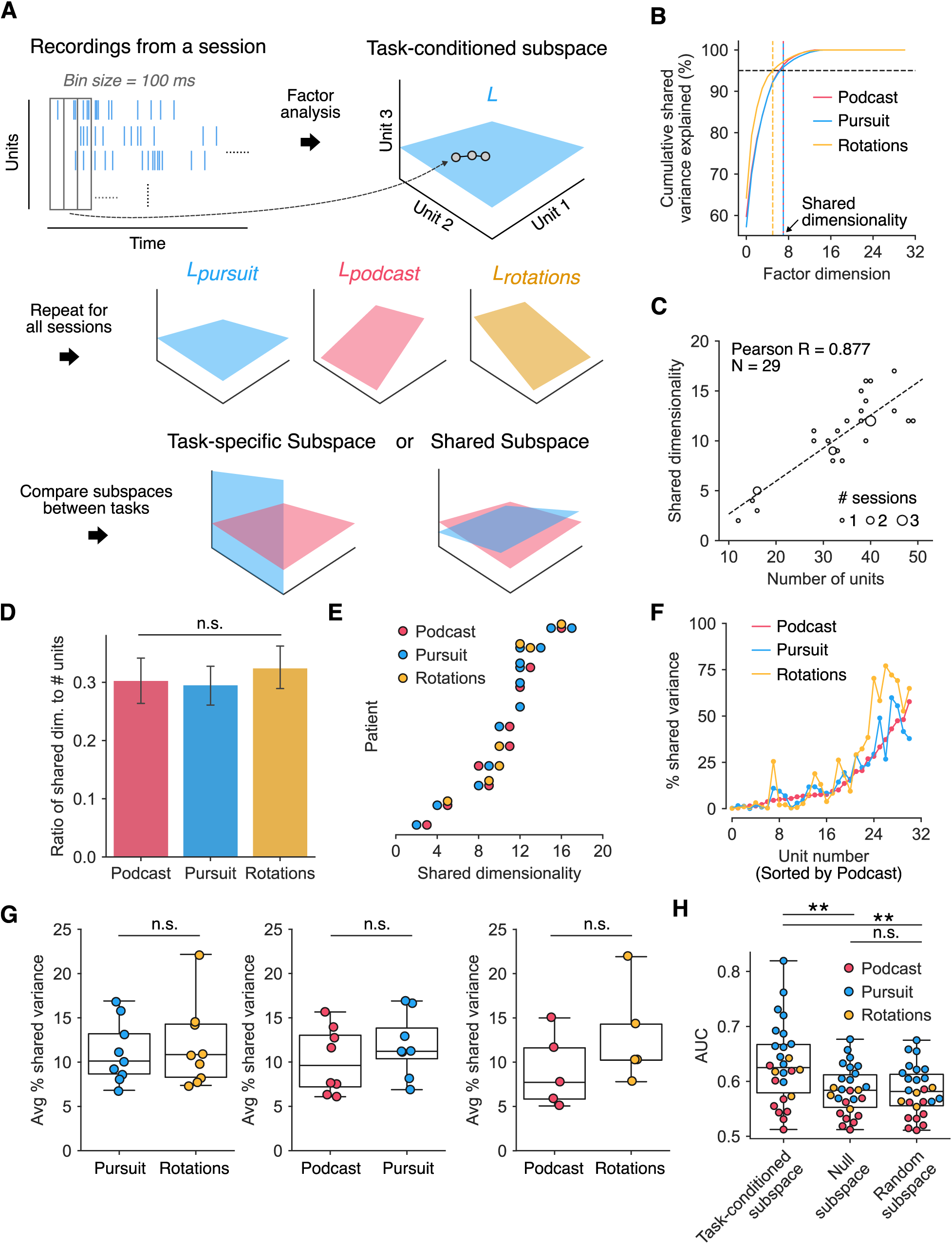
Neural subspaces identified during each task exhibit similar dimensionality and shared variability. **(A)**. Schematic of procedure for identifying a task-conditioned subspace for each task in each patient. Given a matrix of spike counts recorded during a given task (left), we apply Factor Analysis (FA) to identify a task-conditioned subspace (*L*, right) that captures the top dimensions of covariability among spike counts across units. We then compare all pairs of task-conditioned subspaces to test whether subspaces are task-specific or shared (bottom). **(B)**. Percent cumulative shared variance explained by different numbers of factor dimensions in an example patient. The shared dimensionality (indicated by dashed line) is the smallest number of dimensions explaining ≥ 95% of the cumulative shared variance. **(C)**. Relationships between shared dimensionalities and the number of units across all patients. The samples are size-coded by numbers of sessions to better visualize overlapping samples. **(D)**. The ratio of number of units to shared dimensionalities across all sessions for each task. Each bar represents the mean, and error bars indicate the standard deviation across sessions for each task. **(E)**. Shared dimensionality in all sessions per patient. Each dot is the shared dimensionality from a given session. Each row consists of sessions from the same patient. The rows are sorted by the greatest shared dimensionality per patient. **(F)**. Percent of spike count variance that is shared across other units, as identified by Factor Analysis, from an example patient. The unit indices were sorted by the percent shared variance in the Podcast task. **(G)**. Percent of shared variance averaged across all the units in each pair of tasks. Whiskers show the minimum and maximum of the same group. Wilcoxon signed-rank tests assessing differences in means were not significant (n.s.). **(H)**. Areas under the ROC curve (AUC) for decoders trained to distinguish task status (“on” vs. “off”) using different subspace projections. Null subspaces (“null activity”) were obtained by applying factor analysis to independently shuffled spike counts for each unit. Random subspaces (“random activity”) were generated with the same dimensionality as the FA subspace. Null and random results were averaged over 30 repeats. Whiskers indicate the minimum and maximum values within each group. Asterisks indicate statistical significance (**: *p <* 0.01).

To identify each task-conditioned subspace, we used Factor Analysis (FA). FA partitions spike count variability into variability that is shared across units versus variability that is private to each unit (e.g., independent spiking noise; see Methods). This allowed us to characterize a neural subspace without first averaging neural activity across different task conditions. (Using condition-averaged activity would require defining relevant task variables for each *a priori*, which is not straightforward given the high-dimensional and heterogeneous structure of these tasks.) We applied FA to the matrix of spike counts recorded during the trials of each task from a single session (see Methods). We then identified the number of dimensions (or *factors*) that explained 95% of the cumulative shared variance (Fig 3B), termed the *shared dimensionality* [54]. While one could select the dimensionality using cross-validation, our approach led to a more conservative estimate of subspace dimensionality (Fig S2), allowing for a more rigorous test of our hypotheses. This process provided us with a task-conditioned subspace that captured the majority of the covariability between the spike counts of multiple units for each task (see Methods).

Across sessions, shared dimensionality was strongly and positively correlated with the number of units (Fig 3C, Pearson’s *R* = 0.827, *N* = 29 sessions), as commonly observed [9]. Shared dimensionality was similar across tasks, and we did not find any statistical evidence that shared dimensionality divided by the number of units was significantly different for any one task (Fig 3D, Kruskal-Wallis test, *H*(2) = 0.260*, p* = 0.878). Within patients, shared dimensionalities across tasks were similar (Fig 3E).

Performing distinct tasks may require units to change the degree to which their activity covaries with that of other units. To look for evidence of this, we computed the proportion of each unit’s spike count variability that was shared versus private in a given task (*percent shared variance*; see Methods). For an example patient who performed all three tasks, the percent shared variances of units were similar across all tasks (Fig 3F). Across all patients, the average percent shared variance was similar across pairs of tasks (Fig 3G, Wilcoxon signed-rank test, Pursuit and Rotations: *W* = 19.0, *p* = 0.734; Pursuit and Podcast: *W* = 5.0, *p* = 0.078; Podcast and Rotations: *W* = 0.0, *p* = 0.063). Therefore, we did not find evidence that the amount of shared covariance changed across different tasks.

We next wanted to confirm that the task-conditioned subspaces we identified preserved task-relevant information. To do this we analyzed neural population activity after projecting it onto the task-conditioned subspace (see Methods) and found that projected activity continued to encode task-related information. By contrast, projections to a null subspace or a random subspace (see Methods) contained significantly less information (Fig 3H; One-way ANOVA, *F* (2, 75) = 6.87, *p <* 0.01. Post-hoc Tukey tests: FA and null activities: *T* (50) = 2.93, *p <* 0.01; FA and random activities: *T* (50) = 2.96, *p <* 0.01; null and random activity: *T* (50) = 0.08, *p* = 0.935). These results indicate that FA identified a meaningful low-dimensional representation of population activity that preserved task-relevant information.

### Neural subspaces reflect covariability both within and across brain regions

In the previous analyses, we used FA to summarize the activity from both HPC and ACC, treating units without regard to their regional identity. At the single neuron level, we found that HPC units had higher percent shared variances (Fig 4A; two-sided Mann-Whitney U test, *U* = 122312, *p <* 0.01) and higher firing rates (Fig 4B; two-sided Mann-Whitney U test, *U* = 124032, *p <* 0.001) than ACC units. However, it remained unclear whether individual subspace dimensions reflected contributions of one region or both, and whether covariability was predominantly within or across regions. Addressing these questions would clarify how these regions interact during task performance.

**Fig 4.**
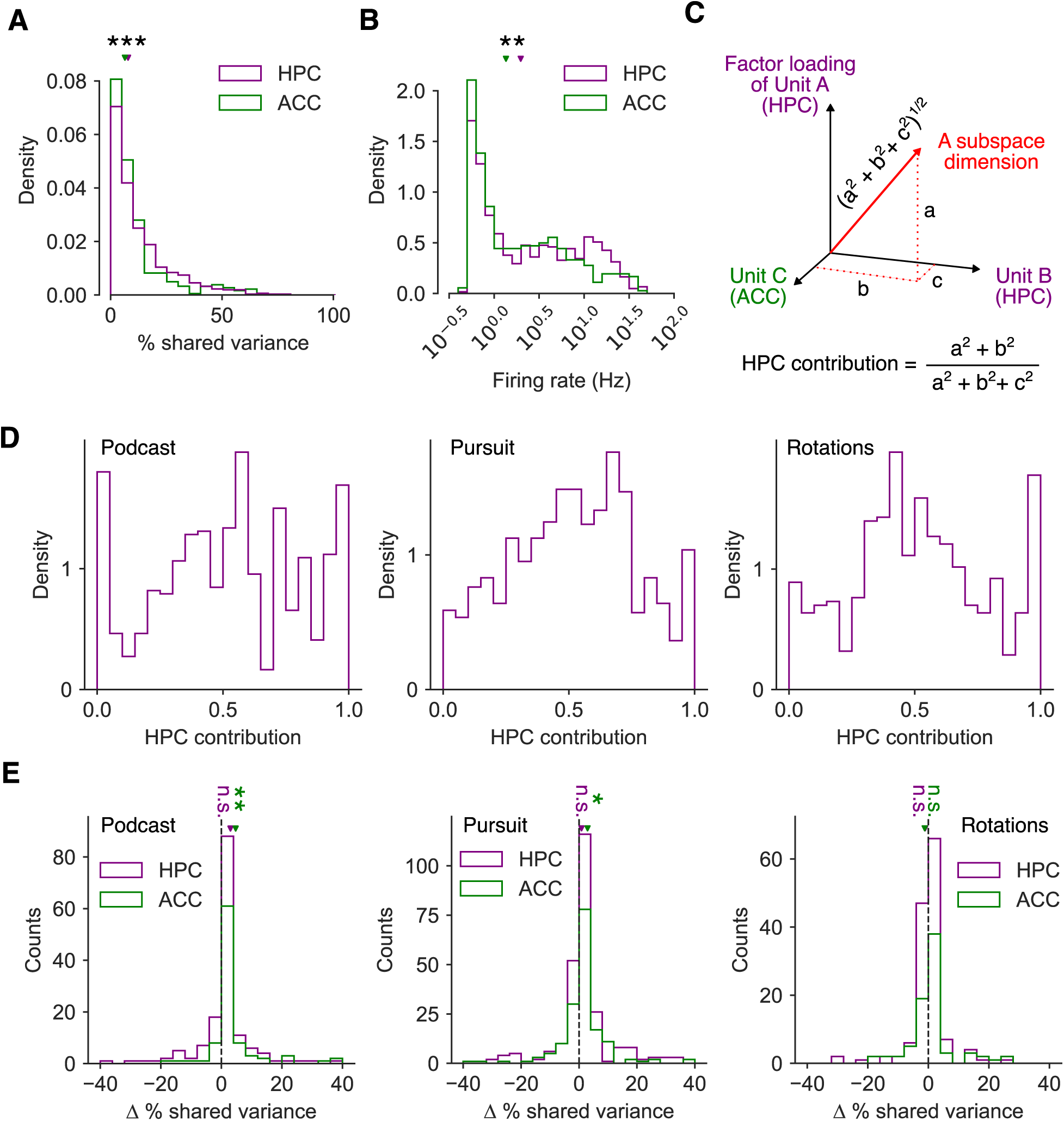
Neural subspaces captured covariability both within and across HPC and ACC units. **(A)**. Distributions of shared variances per unit in individual regions across patients and tasks. Triangles indicate the median of each corresponding distribution. Two-sided Mann–Whitney U tests were used to compare the distributions of shared variance between HPC and ACC units. **(B)**. Distributions of firing rates in individual regions across patients and tasks. Two-sided Mann–Whitney U tests were used to compare the distributions of firing rates between HPC and ACC units. **(C)**. Schematic of estimating each region’s contribution to a subspace dimensions. In this example, there are three units. Red arrow is a subspace dimension with factor loadings of *a*, *b*, and *c* from units A, B, and C, respectively. **(D)**. Subspace contributions from HPC across all dimensions in each task. **(E)**. Distributions of shared variances when including both regions versus when using a single region. Asterisks indicate statistical significance (*: *p <* 0.05; **: *p <* 0.01); n.s., not significant.

We first asked whether subspace dimensions were made up of single or both regions. We hypothesized that most subspace dimensions would be dominated by one region or the other. To test this, we quantified the contribution of each region to each subspace dimension (Fig 4C; see Methods). To control for differences in the number of units from each region, we resampled units within each session so that both regions had the same number of units. Contrary to our hypothesis, we found that subspace dimensions were not exclusively dominated by either HPC or ACC (Fig 4D; 41.3%, 43.4%, and 49.4% of dimensions fell in the center range for Podcast, Pursuit, and Rotations, respectively, whereas 36.6%, 27.9%, and 32.9% fell near the edges). Instead, we found a mixed result: approximately 40% of dimensions fell in an intermediate range, while only around 30% were of dimensions reflected exclusively one region (see Methods). Consistent with this, we found evidence against unimodality for the Podcast and Rotations tasks, but not for the Pursuit task (Fig 4D; Hartigan’s dip test, Podcast: *D* = 0.034*, p <* 0.001; Rotations: *D* = 0.027*, p <* 0.01; Pursuit: *D* = 0.011*, p* = 0.416.) Together, these results indicate that subspace dimensions often reflected contributions from both HPC and ACC, rather than being driven by a single region.

The previous analysis showed that some subspace dimensions included contributions from both regions, suggesting the presence of covariability across brain regions. This implies that estimating subspaces within each region separately may underestimate the amount of variability shared across units. To verify this, we compared the amount of shared variance of each unit when applying FA only to activity from units in the same region, versus when using activity from units in both regions (see Methods). We hypothesized that including the other region would increase percent shared variance. Consistent with this prediction, ACC units showed a significant increase in percent shared variance when HPC units were included in the Podcast and Pursuit tasks (Fig 4E; one-tailed one-sample t-test, Podcast: *T* (98) = 2.90, *p <* 0.01; Pursuit: *T* (177) = 2.06, *p <* 0.05; Rotations: *T* (83) = −0.65, *p* = 0.740). In contrast, we did not observe a corresponding increase for HPC units when ACC units were added (Fig 4E; one-tailed one-sample t-test, Podcast: *T* (184) = 1.55, *p* = 0.061; Pursuit: *T* (278) = 0.71, *p* = 0.239; Rotations: *T* (145) = −0.77, *p* = 0.779). One possible explanation is the imbalance in the number of recorded units across regions, as we had substantially more HPC than ACC units. As a result, adding the smaller ACC population may have had a limited effect on the shared variance structure estimated for HPC units.

Overall, our results suggest that task-conditioned subspaces reflect contributions from both HPC and ACC units. This suggests that interactions between regions were an important component of population structure during these tasks that was best captured when analyzing activity from both regions together.

### Neural subspaces were largely unchanged across tasks

Having identified a neural subspace for each task, we next asked how similar the resulting subspaces were across tasks. Each task-conditioned subspace is a collection of multiple orthogonal dimensions in the space defined by the spike counts of each unit (e.g., as depicted by the two colored lines bordering a given subspace in Fig 1A,B), with coefficients specifying how much each unit contributes to that dimension (e.g., Fig 4C). The factor dimensions are sorted so that the first factor dimension explains the most shared variance, while the last factor dimension explains the least shared variance. For an example patient that accomplished all three tasks, we observed that the first factor dimensions appeared to be similar across all tasks (Fig S3A). To quantify this similarity, we can compute the angles between two of these dimensions. Taking the Podcast and Pursuit tasks in Fig S3A as an example, the angle between the first factor dimensions during the Pursuit and Podcast sessions was 14.90^◦^, substantially lower than what was expected of randomly selected vectors with the same dimensionality (87^◦^ ± 15^◦^, mean ± s.d.. One-tailed t-test, *p <* 0.001, *N* = 500), but similar to the angle between the first factor dimensions of data obtained from the same tasks (14^◦^ ± 23^◦^, mean ± s.d., One-tailed t-test, *p* = 0.223, *N* = 1000, see Methods). This suggests that the first factor dimensions in each task were similar, in tentative support of the Shared Subspace hypothesis (Fig 1B).

In the above analysis, we compared the angle between a single dimension selected from each task-conditioned subspace. However, ideally we would like to know whether the *entire* task-conditioned subspaces, which are collections of multiple orthogonal dimensions, are similar. To do this, we found the principal angles between each task-conditioned subspace [54] (Fig 3A, bottom). Principal angles are a generalization of the notion of “angle” to multiple dimensions: if two subspaces entirely overlap along *K* dimensions, the first *K* principal angles will be zero; if two subspaces are completely orthogonal, all principal angles will be 90^◦^. Because randomly sampled subspaces in high-dimensional space may overlap by chance, we also quantified an upper bound given by the average principal angles expected by randomly sampling subspaces of the same dimensionality (“Random”). Additionally, because non-zero principal angles may occur even when estimating subspaces from the same task (e.g., due to measurement noise), we estimated a lower bound on principal angles by finding subspaces on surrogate sessions constructed by shuffling time steps across both tasks (“Shuffled”; see Methods).

We first visualized the principal angles between an example pair of tasks (Fig 5A). The lowest principal angles were near zero, indicating these task-conditioned subspaces were highly aligned in the first few dimensions. In fact, the principal angles were very close to our lower bound found within the same sessions, indicating that these subspaces were nearly maximally aligned. Across sessions, the principal angles from the data were consistently closer to the lower bound than to the upper bound (Fig 5B), particularly for the first half of the principal angles. To summarize the total distance between two subspaces, we used the geodesic distance (which we will refer to as the *subspace distance*) between pairs of subspaces by taking the *L*2 norm of the principal angles. Using the first half of the principal angles, we found that the subspace distances from each pair of tasks were significantly smaller than random subspaces and were indistinguishable from subspaces within the same tasks (Fig 5C; Podcast and Pursuit: *F* (2, 21) = 68.68, *p <* 0.001; ‘Data’ and ‘Random’: *T* (14) = −8.23, *p <* 0.001, ‘Data’ and ‘Shuffled’: *T* (14) = 2.28, *p* = 0.116. Podcast and Rotations: *F* (2, 12) = 18.09, *p <* 0.001; ‘Data’ and ‘Random’: *T* (8) = −4.25, *p <* 0.01, ‘Data’ and ‘Shuffled’: *T* (8) = 1.25, *p* = 0.735. Pursuit and Rotations: *F* (2, 24) = 51.04, *p <* 0.001; ‘Data’ and ‘Random’: *T* (16) = −7.31, *p <* 0.001, ‘Data’ and ‘Shuffled’: *T* (16) = 2.07, *p* = 0.165.). On the other hand, we did not find evidence that the subspace distances using the second half of the principal angles were significantly different from random or shuffled subspaces (Fig S4A) or in full dimensions (Fig S4B). Together, these results suggest that half of the neural subspace dimensions were highly overlapping across tasks, in support of the Shared Subspace hypothesis.

**Fig 5.**
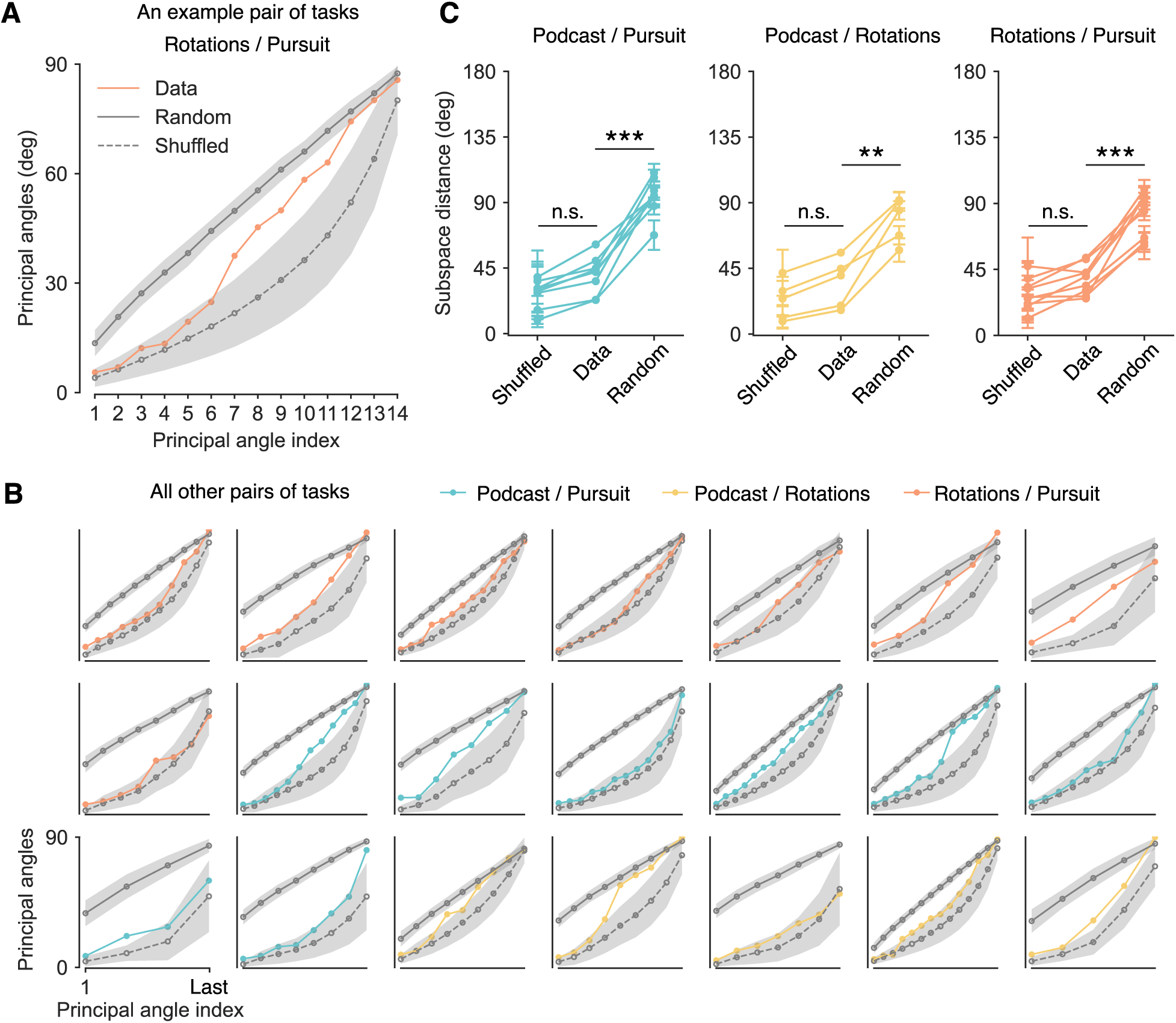
Neural subspaces are highly aligned between tasks. **(A)**. Principal angles from an example pair of tasks. Upper gray line (“Random”) shows average principal angles from randomly estimated subspaces. Lower dashed gray line (“Shuffled”) shows principal angles computed after shuffling time steps across sessions. Shaded areas are standard deviations across 100 repeats. **(B)**. Same as panel A, for all other pairs of tasks. **(C)**. Subspace distances from the first halves of the principal angles in each pair of tasks. Error bars are standard deviations across 100 repeats. One-way ANOVA was followed by pairwise post-hoc Tukey tests with Bonferroni corrections. Asterisks indicate statistical significance (**: *p* < 0.01; ***: *p <* 0.001); n.s., not significant.

### Neural subspaces were aligned in the dimensions containing most neural covariability

Above we showed that the first half of principal angles between neural subspaces were highly overlapping, in support of the Shared Subspace hypothesis (Fig 1B). Next, we asked *where* these subspaces were most aligned. Note that neural variability will be high in some dimensions and low in other dimensions (e.g., factors 1 and 2 explain most of the shared variance, while the remaining factors explain very little variance; see Fig 3B-C). By contrast, our principal angle analysis identifies subspace dimensions with low principal angles regardless of how much neural variance was actually in those dimensions. This raises the question of how the alignment between task-conditioned subspaces relates to the distribution of shared variance across dimensions. We considered three possibilities (Fig 6A). First, task-conditioned subspaces may be aligned only in low-variance dimensions of the population activity (Fig 6A, Case 1). This would suggest that, though the task-conditioned subspaces may overlap, this overlap was largely in dimensions with little neural variance. Second, task-conditioned subspaces may be aligned in high-variance dimensions of the population activity (Fig 6A, Case 2). This would be even stronger support of the Shared Subspace hypothesis, as the dimensions where the task-conditioned subspaces were most aligned would be precisely the dimensions with most of the neural variance. Finally, task-conditioned subspaces may show an inconsistent alignment with low- and high-variance dimensions of the population activity (Fig 6A, Case 3), suggesting that the alignment of task-conditioned subspaces was uncoupled with the distribution of neural variability.

**Fig 6.**
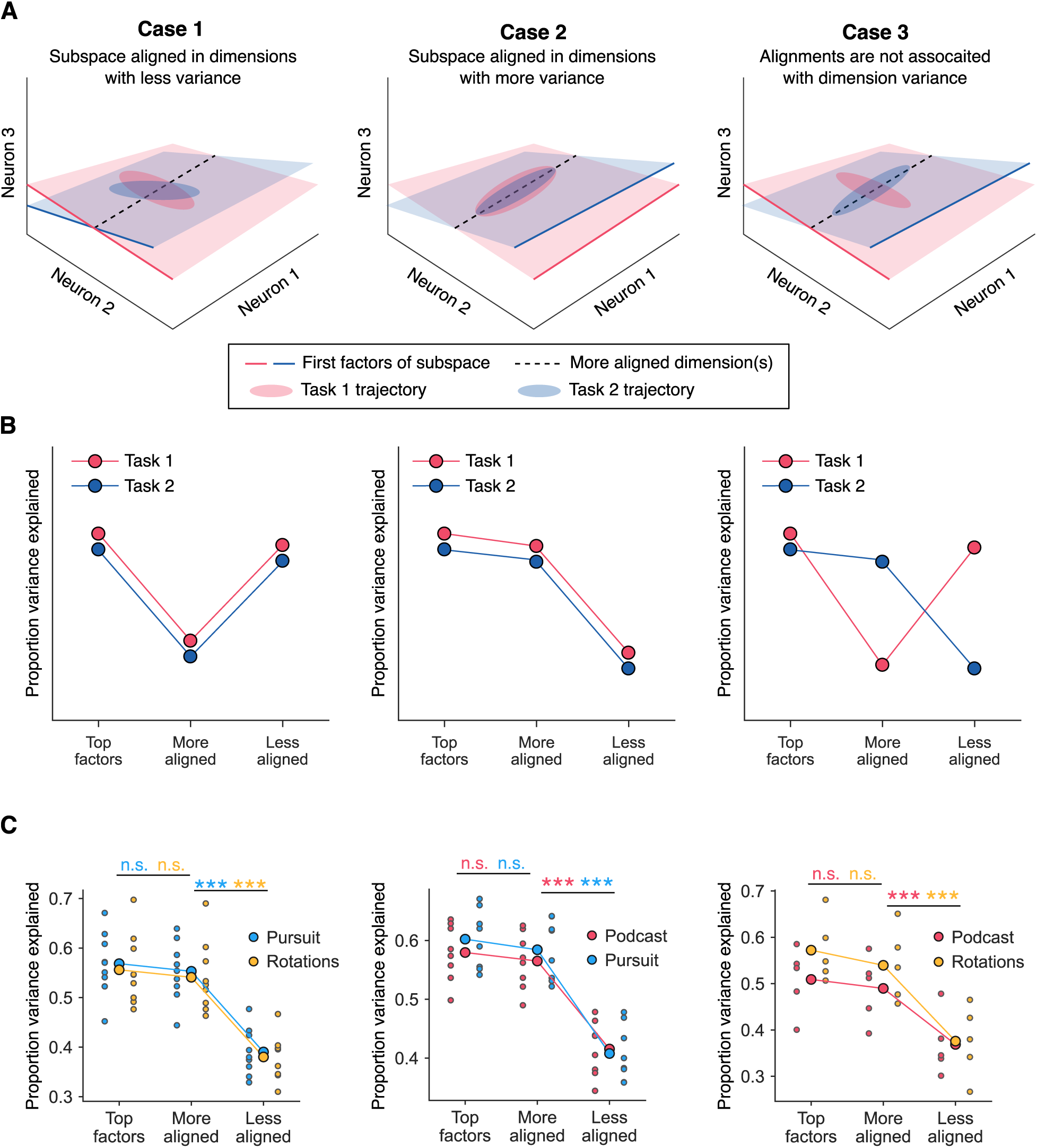
Neural subspaces identified during each task are aligned in dimensions with the most neural variance. **(A)**. Schematic of three hypotheses about how two task-conditioned subspaces (red and blue planes) align with the task-conditioned variance of neural data (red and blue ellipses). The most aligned dimension, indicated by the black dashed line, is the dimension from each subspace with the lowest principal angle. In these cartoons with three neurons, the most aligned dimension of two 2D subspaces is the intersection of the two subspaces. Factor 1 from each subspace (red and blue solid lines) is the dimension within each task-conditioned subspace along which the neural data has the most variance. **(B)**. Illustrations of how proportion explained variance would distribute in different cases. In each panel, the red and blue dots correspond to the red and blue ellipses, respectively. **(C)**. Proportion variance explained by the top *K* factors, the *K* dimensions with the lowest principal angles, and the *K* dimensions with the largest principal angles, using Pursuit/Rotations as example. Big solid dots denote mean across patients, while small dots on both sides of the big dots are from individual patients. Statistical significances are shown for each pair of conditions for different tasks. Asterisks indicate statistical significance (***: *p <* 0.001); n.s., not significant.

To tease apart these possibilities, we performed the following procedure. For each pair of task-conditioned subspaces, we defined a ‘more aligned’ and ‘less aligned’ subspace for each task; these were the subspaces with the *K* lowest and highest principal angles, respectively, where *K* depended on the number of units recorded from each patient (see Methods). We then calculated the percent variance of neural activity in the more aligned and less aligned subspaces from each task, and compared this to the percent variance in the top *K* factors, which contains the maximum amount of variance by construction. By comparing the variance explained in these three categories (i.e., top factors, more aligned, and less aligned), we can delineate where the subspaces align. If the subspaces aligned only in low-variance dimensions (Case 1), the more aligned dimensions would explain the least amount of variance (Fig 6B, left). If they aligned in high-variance dimensions (Case 2), the more aligned dimensions would explain more variance than the less aligned dimensions (Fig 6B, center). And if alignment was inconsistent, the variance explained in those dimensions would be inconsistent as well (Fig 6B, right).

We found that, in nearly every patient and pair of tasks, the more aligned components contained substantially more variance than the less aligned components, and nearly as much variance as the top *K* factors (Fig 6C; Pursuit/Rotations, two-way ANOVA, main effect of categories, *F* (2, 48) = 47.14, *p <* 0.001; Bonferroni-corrected post hoc tests: Top factors vs. Less aligned, *T* (34) = 9.40, *p <* 0.001; More aligned vs. Less aligned, *T* (34) = 8.74, *p <* 0.001; Top factors vs. More aligned, *T* (34) = 0.70, *p* = 0.486. Podcast/Pursuit, two-way ANOVA, main effect of categories, *F* (2, 42) = 66.92, *p <* 0.001; Bonferroni-corrected post hoc tests: Top factors vs. Less aligned, *T* (30) = 10.73, *p <* 0.001; More aligned vs. Less aligned, *T* (30) = 9.79, *p <* 0.001; Top factors vs. More aligned, *T* (30) = 0.94, *p* = 0.353. Podcast/Rotations, two-way ANOVA, main effect of categories, *F* (2, 24) = 15.62, *p <* 0.001; Bonferroni-corrected post hoc tests: Top factors vs. Less aligned, *T* (18) = 5.28, *p <* 0.001; More aligned vs. Less aligned, *T* (18) = 4.43, *p <* 0.001; Top factors vs. More aligned, *T* (18) = 0.78, *p* = 0.446.). These results indicate that the overlap between task-conditioned subspaces occurred predominantly in dimensions with most of the shared variance in neural population activity (as in Case 2 of Fig 6A), in strong agreement with the Shared Subspace hypothesis.

Together, these results provide strong support for the Shared Subspace hypothesis, as task-conditioned subspaces were highly aligned in precisely the dimensions containing most of the neural covariability. This suggests that HPC and ACC population activity largely occupied a consistent low-dimensional subspace regardless of the task being performed.

## Discussion

We recorded neural population activity in human HPC and ACC as patients performed three distinct tasks (Fig 2). We identified a low-dimensional subspace during each task session that captured the dominant modes of neural covariability (Fig 3). We found that, even though neural activity was related to each task individually (Fig 2), this subspace was largely unchanged across tasks (Fig 5, Fig 6), indicating that the correlation structure between neurons remained largely the same despite the distinct cognitive demands of each task. Our results suggest that human HPC and ACC population activity is mostly confined to a shared subspace within the timescale of these experiments (around one day). These findings suggest that neural circuits in HPC and ACC support diverse cognitive functions using a shared subspace.

In this study, we considered three tasks with distinct sensory, cognitive, and motor demands: a natural-language listening task (“Podcast”), a continuous prey-pursuit task (“Pursuit”), and a decision-making task involving mental rotation (“Rotations”). HPC and ACC have been implicated in numerous functions spanning spatial navigation, cognitive mapping, episodic memory, and decision-making, all of which are relevant to the tasks we consider here. Though our analyses here focused on neural covariability rather than tuning to task variables, our results complement recent work that investigated neural tuning in the same data sets. In particular, Chericoni et al. (2025) found that during the Pursuit task, human HPC units encoded the positions of self, prey, and predators [26]; Fine et al. (2025) demonstrated that HPC and ACC units contributed complementary computations to the continuous decision about which prey to pursue [52]; and Franch et al. (2025) revealed that the HPC population carried semantic representations during the Podcast task [53]. Together, these findings highlight the richness of task-encoding in HPC and ACC across three distinct tasks. Our results suggest that HPC and ACC may perform these diverse task-specific computations within a stable, low-dimensional subspace that persists across tasks. Across regions, we found that both regions contributed to the subspace similarly and that covariability was present both within and across regions (Fig 4). Future work is needed to link these findings to the functions of HPC and ACC. Overall, understanding how diverse tasks are implemented within this shared subspace may ultimately reveal general principles of brain organization, with implications for both cognitive theory and clinical interventions. Our findings differ from previous results in mice, where task-specific neurons were observed in parietal cortex [33] and in recurrent neural network models trained to perform multiple tasks, where networks may rely on distinct subspaces to execute multiple tasks with distinct computations [35, 37]. This may be due to differences in our design: the patients in our studies performed our tasks given only minutes of practice, whereas studies of multi-task representations in animals and neural network models typically involve extensive training. We speculate that shared subspaces may be may be more likely during short-term learning, whereas task-specific subspaces may require extended training. Support for this idea comes from studies of primary motor cortex (M1) during learning of different brain-computer interface tasks, where M1 population activity was largely constrained to an *intrinsic manifold* (or shared subspace) during short-term learning [43, 46], and signs of off-manifold activity occurred only following weeks of practice [55]. Such a constraint on neural activity during short-term learning may have computational benefits: As has been proposed in studies of task-switching [56], shared representations may facilitate the rapid acquisition of novel tasks, whereas task-specific representations emerge only with extensive practice.

Our results raise a critical question: how can a shared subspace support flexible, task-specific computations? Flexible computation is often thought to emerge from population-level structure and dynamics rather than from single neurons alone [6, 31, 57–59]. We emphasize that we estimated each task-conditioned subspace using all spiking activity collected during the active trial periods of each session, as opposed to estimating a subspace using the *average* spiking activity in different task conditions. This approach means that each subspace describes the structure of *total* neural covariability, including covariability due to different task conditions and task epochs, but also the covariability due to attentional fluctuations, cognitive strategies, or other internal states. Precisely where task-specific computations occur relative to the shared subspace is not yet known and is an important question for future studies. We consider two possibilities below.

First, task-specific computations may occur *outside* the shared subspace. After all, some subspace dimensions were more strongly shared than others (Fig 5A-B), raising the possibility that the remaining unshared dimensions may indicate the presence of task-specific subspaces. One point against this possibility is that decoding task status was better using activity on versus off the task-conditioned subspace (Fig 3H). Our results also suggest that the unshared dimensions reflect a very small amount of the observed neural variability, as the shared dimensions accounted for the majority of neural variability (Fig 6). This is not totally unreasonable given that spiking activity is known to exhibit large amounts of trial-to-trial variability [60–63]. Because the shared subspace captures total neural covariability, one possibility is that the shared subspace largely reflects task-general information, such as internal states, arousal, and task engagement, while task-specific information is stored only in lower variance dimensions [64].

Alternatively, task-specific computations may occur *within* the shared subspace. For example, because our subspace captures neural variability spanning multiple task epochs, we cannot rule out that multiple “epoch-specific” subspaces are embedded within a shared sub-space (e.g., as in primary motor cortex during preparation versus execution [65, 66] or as in neural networks across task epochs [48]). What would be surprising about this explanation is that, to describe our results, these same subspaces would need to be reused across entirely distinct tasks. There is some evidence for this idea, with work showing that primary motor cortex occupies the same subspace across distinct reaching tasks [16], and that prefrontal cortex and hippocampus reuse overlapping neural populations across different task epochs [44] or even tasks [25, 34, 67]. If the shared subspace indeed contains task-relevant computations across multiple tasks, this suggests the shared subspace acts as a computational “workspace” that can be used as needed to perform different task-specific computations [45,68]. This view is consistent with the “flexibility-via-subspace” hypothesis [47], where task-specific computations occur within a common space defined by low-dimensional factors, rather than neurons. Another related idea is that the brain employs object-centric representations, termed *slots*, that can be flexibly reused and reconfigured across diverse task epochs, enabling rapid generalization and dynamic cognitive mapping [69, 70]. One interpretation of our results in this context is that a shared subspace may arise due to an area reusing the same “slots” across unrelated tasks.

Our study provides suggestive evidence for the possibility that neural populations can process information across distinct sensory, cognitive, and motor domains using a shared subspace. Rather than segregating representations into task-specific structures, the human brain may learn new tasks by relying on a common representational scaffold that can be flexibly reused as task demands change. Such an architecture could underlie the remarkable human capacity for rapid learning and generalization, and it highlights an organizational principle that is largely absent from current neural network models. Looking forward, future work will be needed to test how shared subspaces evolve with long-term training, how they interact with task-specific signal subspaces, and whether similar organizational patterns are observed in other brain regions or disrupted in clinical populations.

## Methods

### Human intracranial neurophysiology

All experiments were performed in the Epilepsy Monitoring Unit (EMU) at Baylor St. Luke’s Medical Center (Houston, Texas, USA). Experimental data was recorded from 11 adult patients (six males and five females) undergoing intracranial monitoring for refractory epilepsy. Neural activity was recorded from Behnke-Fried configuration hybrid macro-micro electrodes (AdTech Medical Instrument Corporation, WI, USA). Each patient had an average of three probes terminating in the left and right HPC, and a similar number in the left and right ACC. Probe locations were verified by co-registered pre-operative MRI and post-operative CT scans. Each probe included eight microwires specifically designed for recording single-neuron activity. All patients had at least one probe terminating in the HPC region, while nine out of 11 patients had at least one probe terminating in the ACC region. Neuron data was recorded using a 512-channel Blackrock Microsystems Neuroport system sampled at 30 kHz.

### Task descriptions

#### Prey-pursuit task (“Pursuit”)

In the Pursuit task [26], patients played 100 trials of a prey-pursuit task. At the beginning of each trial, two shapes, representing an avatar and prey, appeared on a gray background on a standard computer monitor placed in front of the patient. Patients controlled the position of the avatar (a yellow circle, 15-pixels in diameter) on the screen using a joystick (Logitech Extreme Pro 3D). The avatar’s position was limited by the screen boundaries. The prey’s position (square shape, 30 pixels in length) was determined by a simple artificial intelligence algorithm. Each trial ended with either the avatar’s successful capture of the prey or after 20 sec, whichever came first. Successful capture was defined as any spatial overlap between the avatar circle and the prey square. Capture resulted in scored points. In trials with predators, a predator (triangle shape) appeared on 50% of trials. Capture by the predator led to points lost.

### Natural language task (“Podcast”)

During the Podcast task [53] patients listened to six episodes (ranging from five to 13 minutes in duration) taken from The Moth podcast, totaling 47 minutes 25 seconds of listening time (7,346 words). The six stories were “Life Flight”, “The Tiniest Bouquet”, “The One Club”, “Wild Women and Dancing Queens”, “My Father’s Hands” and “Juggling and Jesus”. In each story, a single speaker tells an autobiographical narrative in front of a live audience. The six selected stories were chosen to be both interesting and linguistically rich. Stories were played continuously through the built-in audio speakers of the patient’s hospital television.

### Mental rotations task (“Rotations”)

During the mental rotations task (not previously published), patients performed 100 to 110 trials of a two-alternative forced choice task, where on each trial patients reported whether two 3D shapes presented on a monitor were the same shape or mirror images. One of the two shapes was presented with a different yaw angle, and patients were prompted to mentally rotate the shapes to see if they either matched or were mirrored. The shape pair displayed on each trial was sampled without replacement from one of 384 different options, comprised of 48 possible shapes, 4 different yaw angles for the second shape (0^◦^, 50^◦^, 100^◦^, or 150^◦^), and either mirroring or not mirroring the second shape. Each trial consisted of a variable fixation onset (sampled uniformly between 0.5 and 1.0s), followed by the stimulus onset. Patients then reported their response with an Xbox controller, after which point they were given feedback in the form of a 2-4s video of the shape rotating.

### Neural data analysis

We analyzed neural recordings from 11 human patients while they played a video game (“Pursuit”), listened to podcasts (“Podcast”), or performed a mental rotations task (“Rotations”). We analyzed neural data from a given task session if the patient performed one of the other two tasks within 30 hours. This resulted in 13 sessions of the Pursuit task, 9 sessions of the Podcast task, and 7 sessions of the Rotations task.

### Signal preprocessing

We identified spiking activity by thresholding and denoising the raw voltage traces using the WaveClus algorithm for detecting spikes [71]. To ensure consistency in the neural data across tasks, each session was processed using identical WaveClus parameters with the default WaveClus parameters, except we set the spike detection threshold to 4 times the estimated noise standard deviation (i.e., ‘std_min = 4’) and we only detected negative amplitude deflections (i.e., ‘detection = ‘neg”). Since the same spikes could be detected by multiple channels, we applied the Duplicate Event Removal (DER) algorithm [72]. In brief, DER detects simultaneous spike events across and within electrodes using cross-correlations of recorded units. Afterwards, clusters with abnormal average waveforms or atypical inter-spike intervals were manually removed. The spiking activity in the remaining clusters on the same channel was then treated as a single multi-unit neural source.

For each task, for each channel with detected unit activity, we found the spike counts of *N* different units in *T* = 100 ms nonoverlapping bins. The result was a *N* × *T* matrix of spike counts for a given patient and task. The number of units per patient was 37 ± 14 (mean ± s.d., *N* = 11 patients), while the number of time bins (*T*) varied per task as follows. Pursuit spiking data came from experiments comprised of 100 trials, resulting in 15, 379 ± 4, 529 time bins per session (mean ± s.d., *N* = 11 sessions). Note that two patients completed two Pursuit sessions each. Podcast spiking data came from stories with 7, 341 spoken words, resulting in 26, 165 ± 3, 969 time bins per session/patient (mean ± s.d., *N* = 9 sessions). Rotations spiking data came from 100 (*N* = 5 sessions) or 110 trials (*N* = 2 sessions) of the task, resulting in 7, 359 ± 1, 904 time bins per session/patient (mean ± s.d., *N* = 7 sessions).

### Task-tuning of individual units

For single-neuron tuning analyses, spike trains from representative units were aligned to task-relevant behavioral events within each session and analyzed at millisecond resolution. In Podcast sessions, spikes were aligned to word onset; in Pursuit sessions, spikes were aligned to trial onset; and in Rotations sessions, spikes were aligned to trial offset. For each event, perievent spike segments were extracted using predefined pre- and post-event windows. These spike trains were then convolved with a Gaussian kernel to obtain smoothed estimates of instantaneous firing rate, and event-aligned responses were averaged across trials to generate peri-event time histograms. Variability across trials at each time point was quantified as the standard error of the mean (SEM).

To assess event-related modulation relative to ongoing activity, each neuron’s firing during the peri-event period was compared with its average firing rate across the full recording session. At each peri-event time point, a two-tailed one-sample t-test was used to test whether trial-wise firing rates differed from the session-wide baseline mean. Resulting p values were corrected across time points within each peri-event trace using false discovery rate (FDR) correction.

### Comparing mean firing rates across tasks

We computed mean firing rates for units recorded in at least two tasks within the same patient, using firing rates aggregated across entire sessions. Changes in firing rates were quantified using Cohen’s *d* (unpaired), computed with pooled standard deviation to compare differences in means relative to variability. We used a threshold of *d* = 0.5, a conventional cutoff for moderate effect size, to classify the magnitude of changes in firing rates.

### Factor Analysis

We fit a Factor Analysis (FA) model to the spike count data from each task, for each patient. To ensure the factor analysis captured variability that was shared across all units, we first z-scored each unit’s activity to have zero mean and unit standard deviation, so that units could contribute equally regardless of their firing rates. The FA model is given by:

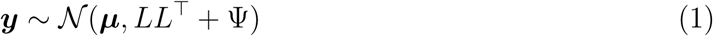

where ***y*** ∈ R*^N^* are the z-scored spike counts from *N* simultaneously recorded units, ***µ*** ∈ R*^N^* is the empirical mean of ***y***, *L* ∈ R*^N^*^×^*^M^* is the loading matrix relating *M* factors to our *N* -dimensional spike counts, and Ψ ∈ R*^N^*^×^*^N^* is a diagonal matrix of independent variances for each unit. We estimated the FA parameters ***µ***, *L*, and Ψ using scikit-learn with svd_method = lapack, and tol = 1*e* − 6.

### Task-conditioned subspace

We now describe how we defined the task-conditioned subspace, the low-dimensional linear subspace that captures dominant patterns of co-modulation across neural units. Because each task had a substantial number of time steps, the dimensionality maximizing the cross-validated log-likelihood was often large (Fig S2). To obtain a more conservative estimate of the dimensionality we estimated the FA parameters using *M* = *N*, and then followed previous work by defining the shared dimensionality, *d_shared_*, as the number of dimensions needed to explain 95% of the cumulative shared covariance (Fig 3B-C) [46, 54]. Here, the shared covariance explained by factor *i* is given by the *i^th^* eigenvalue of *LL*^⊤^. We next defined *L̃* as the first *d_shared_* columns of *L*, so that 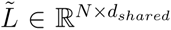. The dimensions defined by the columns of this matrix are stretched by the amount of variance they explain, so we next found an orthonormal basis for the columns of *L̃* using singular value decomposition [73]. We defined the task-conditioned subspace as the resulting orthonormal basis for *L̃*, which we will refer to for simplicity as *L*.

We calculated the percent shared variance, or the percentage of each unit’s variance that was shared versus private [46, 54], as follows:

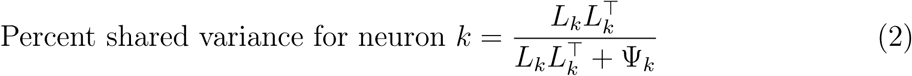

where *L_k_* is the *k^th^* row of *L*, and Ψ*_k_* is the independent (i.e., private) variance of unit *k*.

### Subspace analysis for different brain regions

For each session, we applied FA to the population activity including units from both HPC and ACC. We then computed the percent shared variance for each neuron and grouped these values by brain region (Fig 4A).

To quantify contributions of a region to a subspace (Fig 4D), we summed the squares of factor loadings (*n*) in that regions and normalized it by the total sum of squares. If the task-conditioned subspace was orthonormal, the denominator would reduce to unity. In this study, because we considered only two brain regions, we characterized contributions in terms of HPC, with ACC contributions defined as the complement (i.e., the two contributions sum to one). We then aggregated HPC contributions across all subspace dimensions and sessions within each task.

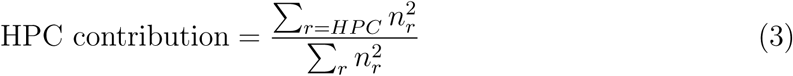

However, since there were a different number of units in each region, this contribution metric is biased toward the region with more units, usually HPC. Therefore, we repeated the process (*N* = 10) with randomly resampled units such that both regions had the same number of units.

To quantify HPC contributions, we defined a “center” range as values between 0.3 and 0.7, and an “edge” range as values less than 0.2 or greater than 0.8. These ranges were chosen to cover equal intervals, ensuring a balanced comparison. Dimensions in the edge range were dominated by either HPC or ACC, whereas dimensions in the center range reflected more balanced contributions from both regions.

To further quantify the shape of the distribution, we used Hartingan’s dip test [74] to quantify if the distribution is unimodal. The null hypothesis of Hartingan’s dip test is unimodality. Therefore, significant values indicate evidence against a unimodal distribution. To quantify cross-regional covariability (Fig 4E), we compared the percent shared variance of each unit when FA was fit to a single region versus when it was fit jointly to both regions. We relied on cross-validated FA rather than the conventional 95% cumulative explained variance rule because the 95% rule can bias factor loadings toward high-variance units. This happens because the method minimizes dimensionality by favoring those units in order to reach the 95% threshold. In contrast, cross-validated FA identifies the subspace that best captures shared covariance regardless of dimensionality, avoiding this bias.

### Pairwise cross-task analysis

To ensure that neural population activity and their corresponding subspaces were directly comparable across tasks, we restricted our analysis to the subset of channels recorded in both sessions. For each task pair, we re-estimated the task-conditioned subspaces using only these mutual channels, applying the same shared-variance thresholding method described above.

### Principal angles analyses

Suppose we have two task-conditioned subspaces, *L*_1_ ∈ R*^N^*^×^*^K^*^1^ and *L*_2_ ∈ R*^N^*^×^*^K^*^2^, with *K*_1_ and *K*_2_ dimensions, and each with orthonormal columns. Let *s*_1_*, . . ., s*_min(*K*_ *_,K_* _)_ ∈ R^+^ be the singular values of 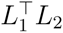. Then the *k^th^* principal angle (in radians), *θ_k_*, is given by *θ_k_* = cos^−1^(*s_k_*). We can then calculate the total distance (in radians) between these subspaces using the geodesic distance [75], calculated as:

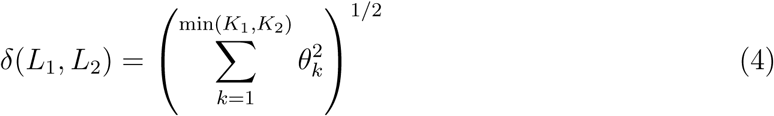

To quantify the chance level (i.e., the upper bound) of principal angles in two task-conditioned subspaces, *L*_1_ ∈ R*^N^*^×^*^K^*^1^ and *L*_2_ ∈ R*^N^*^×^*^K^*^2^, we first sampled 100 random orthonormal matrices *L̂*_1_ ∈ R*^N^*^×^*^K^*^1^ and *L̂*_2_ ∈ R*^N^*^×^*^K^*^2^ . Next, we simulated neural activity ***ŷ***_1_ and ***ŷ***_2_ by:

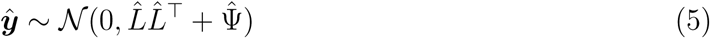

where Ψ̂ = 1*e*^−4^. We then estimated subspaces using ***ŷ***_1_ and ***ŷ***_2_ with specified shared dimensionalities *K*_1_ and *K*_2_, and we computed principal angles using the above pipeline.

To obtain the lower bounds of principal angles, we should estimate the subspaces in the same data. Therefore, we randomly partitioned the neural population activity ***y*** ∈ R*^T^*^×^*^N^* into 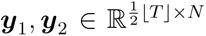. We estimated their neural subspaces, *L*_1_*, L*_2_ with the same shared dimensionality, *K*, as ***y***. We then computed principal angles using the above pipeline.

### Subspace alignment

To calculate the amount of shared variance in the least versus most aligned dimensions of the task-conditioned subspaces (Fig 6D-E), we performed the following procedure for each pair of task-conditioned subspaces, *L*_1_ and *L*_2_. We first found the left and right singular vectors of 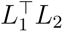. The *i^th^* left and right singular vectors, ***ℓ***_1_(*i*) ∈ R*^N^*, and ***ℓ***_2_(*i*) ∈ R*^N^*, are the dimensions of *L*_1_ and *L*_2_, respectively, corresponding to the *k^th^* principal angle, *θ_k_* (i.e., the angle between ***ℓ***_1_(*i*) and ***ℓ***_2_(*i*) is *θ_k_*). Each task-conditioned subspace has *K*_0_ = min(*K*_1_*, K*_2_) such singular vectors. We then split these *K*_0_ principal vectors into two equal groups, of size 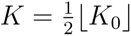, based on whether they corresponded to the *K* smallest or largest principal angles. These two groups of vectors define two subspaces, which we call the “More Aligned” subspace and “Less Aligned” subspaces, respectively. For each task—say, task 1—we then found the amount of shared variance in the More Aligned and Less Aligned subspaces by finding the total variance of the z-scored spike counts recorded during the corresponding task in these two subspaces. We compared this to the total variance in the first *K* columns of the task-conditioned subspace, *L*_1_ (“Top factors” in Fig 6D-E), as this subspace has the most shared variance by construction.

### Decoding analysis

#### Decoding task status using normalized population activity

To quantify task-related structure in population activity, we defined a binary task-status variable for each time bin indicating whether the subject was behaviorally engaged in the task (“on”) or not (“off”). This definition was adapted to the temporal structure of each paradigm. For the Pursuit and Rotations tasks, “on” bins spanned the interval from trial onset to trial offset (Fig 2A,C), whereas for the Podcast task, “on” bins were defined by the presence of auditory word stimuli (Fig 2B). Decoding was performed separately within each recording session.

Task status was decoded from neural population activity using logistic regression with a liblinear solver (Fig 2J). The decoder was trained on normalized spike-count activity, or activity projected into a low-dimensional FA subspace. Decoder performance was evaluated with stratified 10-fold cross-validation, such that each fold preserved the proportion of “on” and “off” bins. Within each fold, features were standardized using the training data only, and the same transformation was then applied to the held-out data to avoid information leakage. Because behavioral engagement was not necessarily balanced across bins, decoder performance was quantified primarily using the area under the ROC AUC, which provides a threshold-independent measure of separability under class imbalance.

### Decoding task status using subspace activity

Subspace activity decoding (Fig 3H) was performed separately within each recording session to test whether population activity distinguished behavioral on versus off periods. Binary labels were defined from the session’s on/off annotations, and decoding was carried out independently for several feature representations. For the empirical latent-space condition, normalized spike-count activity was projected into the task-conditioned subspace estimated for that session, yielding low-dimensional task-conditioned subspace activity. For control conditions, we generated either a null subspace by fitting FA after independently permuting each neuron’s normalized activity across time, or a random orthonormal subspace with dimensionality matched to the session’s FA shared dimensionality.

For each representation, decoding was performed using logistic regression with a liblinear solver under stratified 10-fold cross-validation. Within each fold, features were z-scored using statistics computed from the training partition only, and the same transformation was then applied to the held-out partition. Decoder performance was quantified primarily by held-out ROC AUC. For the randomized control representations, decoding was repeated 30 times per session using independently generated null or random subspaces, and the resulting session-level performance was summarized as the mean across repeats.

To compare decoding performance across subspace conditions, session-level ROC AUC values were aggregated across the three groups: task-conditioned subspace, null subspace, and random subspace. Group differences were assessed using a one-way analysis of variance (ANOVA), followed by Bonferroni-corrected post hoc pairwise comparisons.

### Decoding task identity

To quantify task-related information in neural population activity, we decoded task identity from raw spike-count activity using pairwise classification between tasks (Fig 2K). For each task pair, the analysis was restricted to the subset of channels recorded in both sessions (see Pairwise cross-task analysis), ensuring that decoding was performed on matched neural populations across tasks. Decoding was performed separately for each task pair using logistic regression with a liblinear solver.

Classifier performance was evaluated with stratified 10-fold cross-validation, such that each fold preserved the relative proportion of samples from the two task conditions. Within each fold, features were standardized using the training data only, and the same transformation was then applied to the held-out data to avoid information leakage. Decoder performance was quantified primarily by the ROC AUC, which provided a threshold-independent measure of task discriminability.

### Statistical analysis

All statistical analyses were performed using custom Python code. Unless otherwise stated, tests were two-sided. Depending on the comparison of interest, we used one-sample, paired-sample, and independent-sample t-tests; Wilcoxon signed-rank tests; Wilcoxon rank-sum tests; Kruskal-Wallis tests; Pearson correlation tests; and one-way or repeated-measure ANOVA. When ANOVA effects were significant, post hoc pairwise comparisons were performed with multiple-comparison correction (e.g., Bonferroni correction). Effect sizes were quantified with Cohen’s d where appropriate. Sample sizes were not predetermined by formal statistical methods.

## Acknowledgements

This work was supported by the McNair Foundation and by NIH U01 NS121472.

## Supplementary Figures

**Fig S1.**
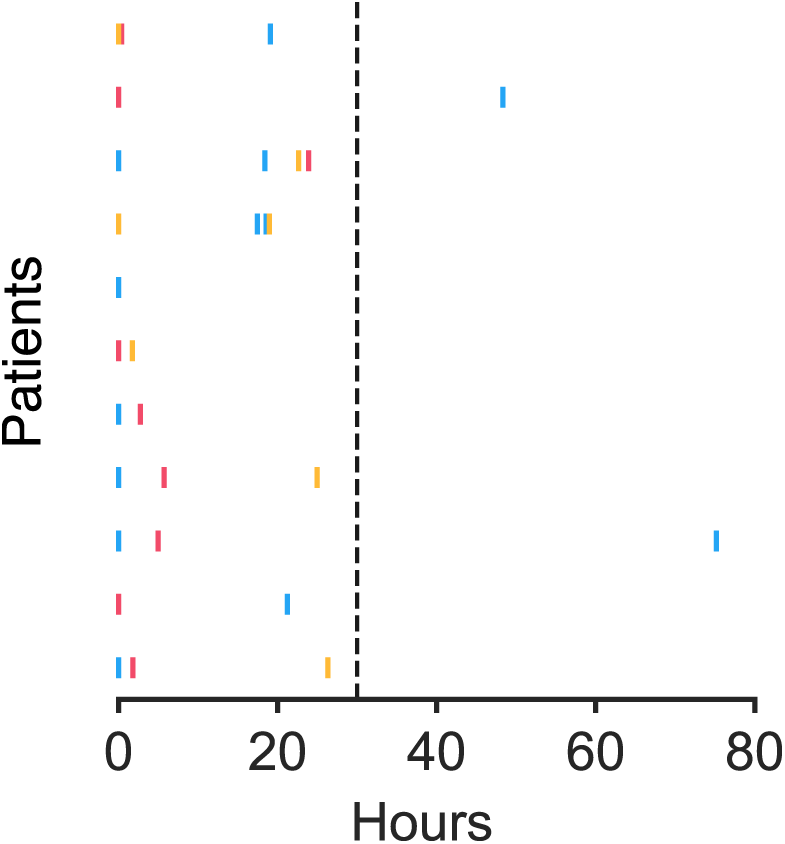
Task timing for all 11 patients, aligned to the time of each patient’s first task. All sessions were included in single-session analyses. For pairwise analyses, we excluded pairs of tasks that were performed more than 30 hours apart (i.e., two Pursuit sessions from Patient C and Y).

**Fig S2.**
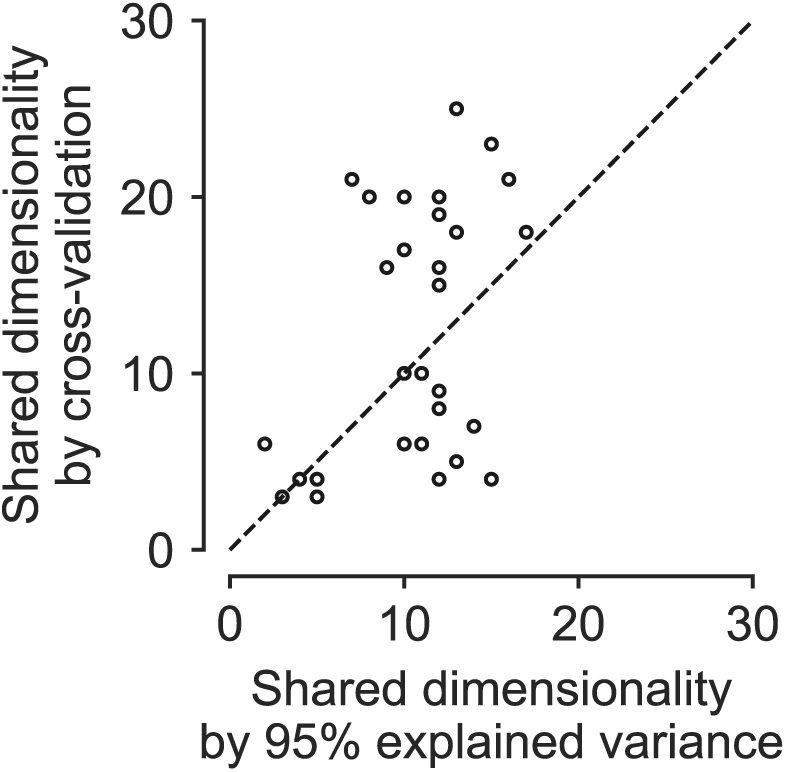
Shared dimensionality obtained from different approaches. Each data point indicates the shared dimensionality in a session estimated by the minimal number of factors that explained 95% cumulative variance and by cross-validation. The diagonal line indicates identical shared dimensionalities.

**Fig S3.**
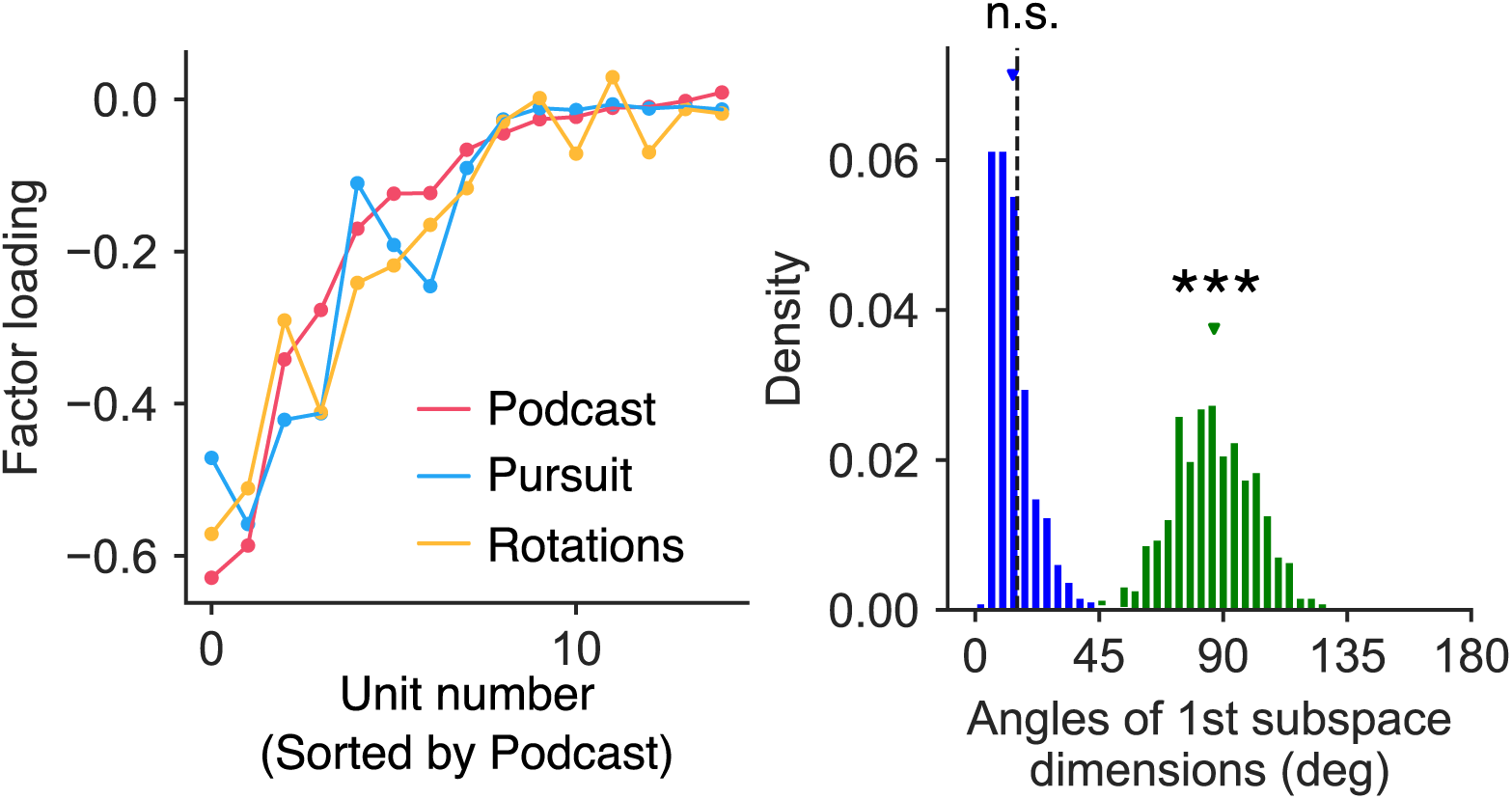
Factor loadings of the first dimensions are similar across tasks. **(A)**. The factor loadings of the first dimensions from a patient that accomplished all three tasks. The unit indices were sorted by the factor loading in the Podcast task. **(B)**. The angle between the first dimensions of the Podcast and Pursuit tasks and their upper and lower bounds, as represented by randomized subspaces (“Random”) and shuffled neural data (“Shuffle”), respectively. Asterisks indicate statistical significance (***: *p <* 0.001); n.s., not significant.

**Fig S4.**
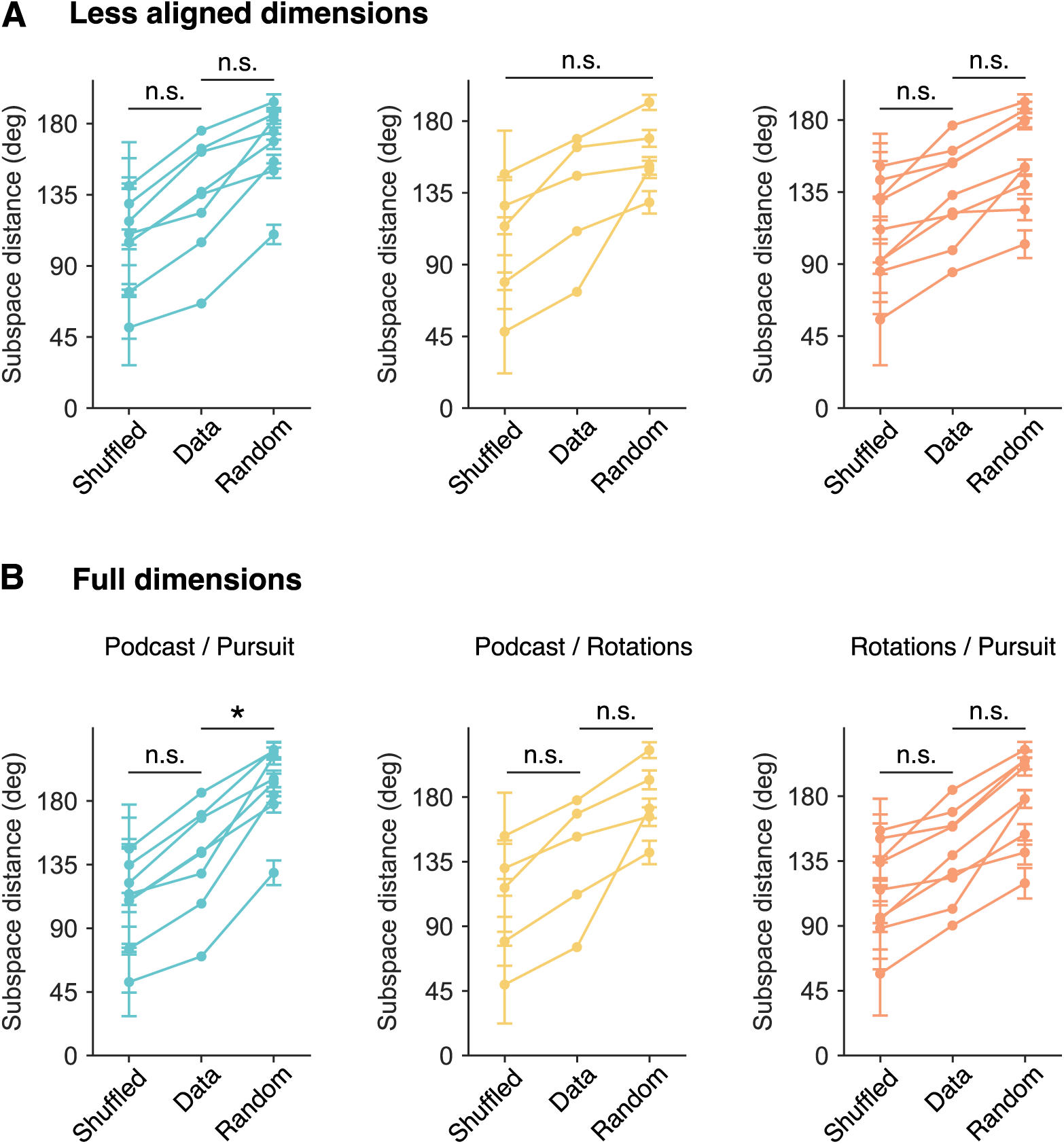
Subspace distance analysis. Similar to Fig 5C, we compare the subspace distances of each pair of tasks with the upper and lower bounds. **(A)**. Subspace distances estimated across less aligned dimensions. Podcast and Pursuit: *F* (2, 21) = 7.86, *p* = 0.003; ‘Data’ and “Random’: *T* (14) = −2.00, *p* = 0.196, ‘Data’ and ‘Shuffled’: *T* (14) = 1.81, *p* = 0.278. Podcast and Rotations: *F* (2, 12) = 3.08, *p* = 0.083. Pursuit and Rotations: *F* (2, 24) = 5.02, *p* = 0.015; ‘Data’ and ‘Random’: *T* (16) = −1.56, *p* = 0.418, ‘Data’ and ‘Shuffled’: *T* (16) = 1.63, *p* = 0.369. **(B)**. Subspace distances estimated across all the dimensions. Podcast and Pursuit: *F* (2, 21) = 13.04, *p <* 0.001; ‘Data’ and ‘Random’: *T* (14) = −3.03, *p* = 0.027, ‘Data’ and ‘Shuffled’: *T* (14) = 1.85, *p* = 0.259. Podcast and Rotations: *F* (2, 12) = 4.47, *p* = 0.035; ‘Data’ and ‘Random’: *T* (8) = −1.76, *p* = 0.351, ‘Data’ and ‘Shuffled’: *T* (8) = 1.18, *p* = 0.813. Pursuit and Rotations: *F* (2, 24) = 8.61, *p* = 0.002; ‘Data’ and ‘Random’: *T* (16) = −2.50, *p* = 0.072, ‘Data’ and ‘Shuffled’: *T* (16) = 1.66, *p* = 0.351. Asterisks indicate statistical significance (*: *p <* 0.05); n.s., not significant.

